# Inhibition of Extracellular Vesicle-Associated MMP2 Abrogates Intercellular Transfer of Hepatic miR-122 to Tissue Macrophages and Curtails Liver Inflammation

**DOI:** 10.1101/2021.01.04.425217

**Authors:** Arnab Das, Sudarshana Basu, Diptankar Bandyopadhyay, Debduti Dutta, Sreemoyee Chakrabarti, Moumita Adak, Snehasikta Swarnakar, Partha Chakrabarti, Suvendra N. Bhattacharyya

## Abstract

microRNA-122 (miR-122), a liver specific regulatory RNA, plays an important role in controlling metabolic homeostasis in mammalian liver cells. Interestingly, miR-122 is also a proinflammatory microRNA and when exported to tissue resident macrophage induces expression of inflammatory cytokines there. We found intercellular transfer of miR-122 in lipid exposed liver plays a role in liver inflammation. Exploring the mechanism of intercellular miR-122 transfer from hepatic cells, we detected MMP2 on the membrane of extracellular vesicles derived from hepatic cells which proved to be essential for transfer of extracellular vesicles and their miRNA content from hepatic to non-hepatic cells. Matrix metalloprotease 2 or MMP2 is a metalloproteinase that plays a key role in shaping and remodelling the extracellular matrix of human tissue by targeting degradation of matrix proteins. MMP2 was found to increase the movement of the EVs along the extracellular matrix to enhance their uptake in recipient cells. Inhibition of MMP2 restricts functional transfer of hepatic miRNAs across the hepatic and non-hepatic cell boundaries. By targeting MMP2, we could reduce the innate immune response in mammalian liver by preventing intra-tissue miR-122 transfer.

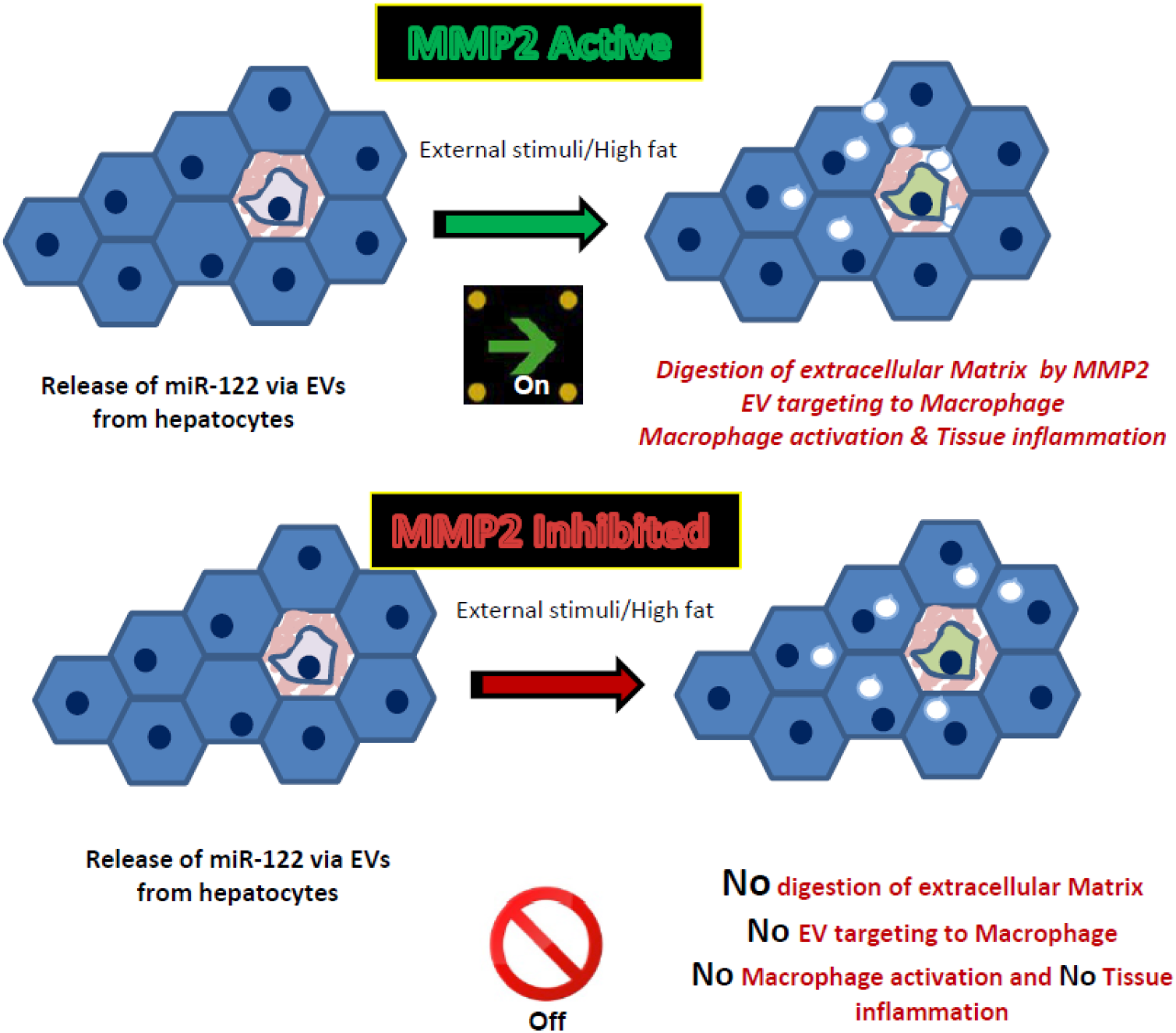

- Human hepatocytes on exposure to high lipid export out miRNAs including proinflammatory miR-122.
- Extracellular miR-122 is taken up by tissue macrophages to get them activated to produce inflammatory cytokines.
- MMP2 present on the surface of the EVs released by hepatocyte is essential for miRNA transfer to macrophage cells
- Inhibition of MMP2 prevents miR-122 transfer to macrophage and stops activation of recipient macrophage.

## Introduction

miRNAs are 22 nt long regulatory RNAs that can repress protein synthesis from their target mRNAs by imperfect base pairing (Bartel, 2018; Filipowicz et al, 2008). Majority of mammalian genes are under miRNA regulation and deregulation of miRNA activity and expression is associated with human diseases (Bartel, 2009). These tiny regulatory RNAs that primarily affect the post-transcriptional steps of gene expression can also be transferred across cell boundaries. Extracellular vesicle (EV) or exosome mediated delivery of miRNA acts as a major way of exchange of genetic information between cells in mammalian tissues and organs (Valadi et al, 2007). The pathology of different diseases is found to be associated with deregulation of miRNA machineries including its export via EVs. Export of miRNAs via EVs thus, not only plays a critical role in maintaining the homeostasis of gene expression in higher eukaryotes but also allows rapid response by recipient cells under stress (Mukherjee et al, 2016).

Hepatic miRNA., miR-122, identified as a highly abundant liver-specific miRNA, has a significant role in controlling hepatic metabolic processes (Chang et al, 2004). Inhibition of miR-122 leads to down-regulation of a number of lipogenic and cholesterol biosynthesis genes in the liver (Esau et al, 2006; Jopling, 2012). This in turn leads to reduced plasma cholesterol, increased hepatic fatty-acid oxidation and reduced synthesis of hepatic lipids. Various studies on circulating miRNAs in NAFLD patients reveal a distinctive serum miRNA profile based on the progression of disease. Upregulated levels of miR-122, miR-192 and miR-375 could be correlated with disease severity in NASH patients as compared to patients with steatosis only (Pirola et al, 2015).

The extracellular vesicles or exosomes are 30-100 nm diameter particles and presence of specific proteins and RNA cargo within the EVs make them unique and cell type specific and thus selectively get delivered to target cells where cell-type specific uptake occurs either by receptor mediated endocytosis or phagocytic activity of the recipient cells.

EV-entrapped miRNAs are also secreted into the extracellular milieu (blood, serum or plasma) and circulating miRNAs, being identified as potential biomarkers for metabolic diseases, can act as paracrine and endocrine signals in circulation, and can influence the target gene expression in recipient cells (Basu & Bhattacharyya, 2014; Wang et al, 2019). Recently, a study showed that in alcohol fed mice, miRNA-122 transferred via Extracellular Vesicles (EVs) to monocytes and liver Kupffer cells sensitized them to LPS stimulation and pro-inflammatory cytokine production (Momen-Heravi et al, 2015).

How the EVs with their cargo get transferred through the extracellular matrix before they get delivered to recipient cells is an interesting question and the matrix metalloproteases are interesting candidates to explore (Shimoda & Khokha, 2017). Recently, MMP2 has been identified as a metalloprotease present in the EV released by osteoblast cells and found to be essential for endothelial cell angiogenesis (Tang et al, 2019). MMP2 has been found to be recruited to membranes via MMP14 that is known to act as an adopter protein of MMP2 for its association with biomembranes (Han et al, 2015). MMP2 in its mature form is an abundant matrix metalloprotease and its level in the matrix is heavily regulated and found to be altered in liver disease context (Wang et al, 2014). In different types of cancers the importance of MMP2 is well documented (Han et al, 2020).

In this manuscript, we have documented the presence of MMP2 in the EVs isolated from hepatic cells. The EV association of MMP2 is MMP14 dependent and we have also noted that the EVs with MMP2 can ensure functional cell to cell miRNA transfer. This process is blocked by the MMP2 inhibitor and restored by recombinant MMP2. The transfer of pro-inflammatory miR-122 from hepatocytes to liver resident macrophage cells is dependent on MMP2. We have also documented the retardation of movement of EVs in collagen matrix by MMP2 inhibitor. Thus MMP2 can facilitate the functional miRNA cargo transfer across cell boundaries via EVs and by blocking the EV-transport with MMP2 inhibitor the expression of inflammatory cytokines can be perturbed in mammalian liver.

## Results

### Transfer of hepatic miR-122 to recruited monocytes and Kuppfer cells in Methionine Choline Deficient (MCD) diet fed mice livers

Obesity is known to be accompanied by a low grade, chronic inflammatory response orchestrated by pro-inflammatory cytokines like IL-1β, TNF-α, IL-6, CCL2 and others (Berg & Scherer, 2005; Shoelson et al, 2006). The liver is also a site for inflammation in the obese state (Cai et al, 2005). Macrophages are known to play a significant role in hepatic inflammation and subsequent development of insulin resistance (Alisi et al, 2017). The hepatic macrophage population is a heterogenous one, composed of resident Kupffer cells (KC) and infiltrating macrophages (Recruited Hepatic Macrophages-RHM). We wanted to explore the importance of liver specific miRNAs in activation of tissue macrophages on exposure to MCD diet. To address this point, we chose to study a mice model of NASH by feeding the mice with MCD diet for 30 days (Caballero et al, 2010) and FACS sorting the two macrophage populations by differentially labelling them with fluorescent markers (Figure 1A). MCD diet fed mice livers showed increased fat accumulation and hepatocyte ballooning characteristic of NASH (Figure 1B). Normal chow (Ch) fed mice livers show none of these characteristics of NASH. Non-Parenchymal cells were isolated from hepatic cell suspensions prepared from both MCD and Chow diet fed mice livers followed by FACS sorting of F4/80^+^/CD11b^+^ dual marker positive KC cells (R5) and F4/80^low^/CD11b^+^ RHM cells that have low expression of F4/80 (R4) (Figure 1A) (Morinaga et al, 2015). The F4/80^low^/CD11b^+^ set (R4) was sorted out as this represents fresh RHMs derived from circulating monocytes (Morinaga et al, 2015). The CD11b^+^ population in MCD diet fed mice livers appeared as two discrete fractions-the F4/80^low^/CD11b^+^ set (R4) and the F4/80^high^/CD11b^+^ set (R8). R8 appeared to be negligible in Ch diet fed mice and probably represents matured differentiated macrophages and was not included in the analysis (Morinaga et al, 2015). We restricted our investigations to the R4 population as they represent fresh RHMs and hence would be more representative of conditions leading to infiltration and inflammation (Morinaga et al, 2015).

**Figure 1.**
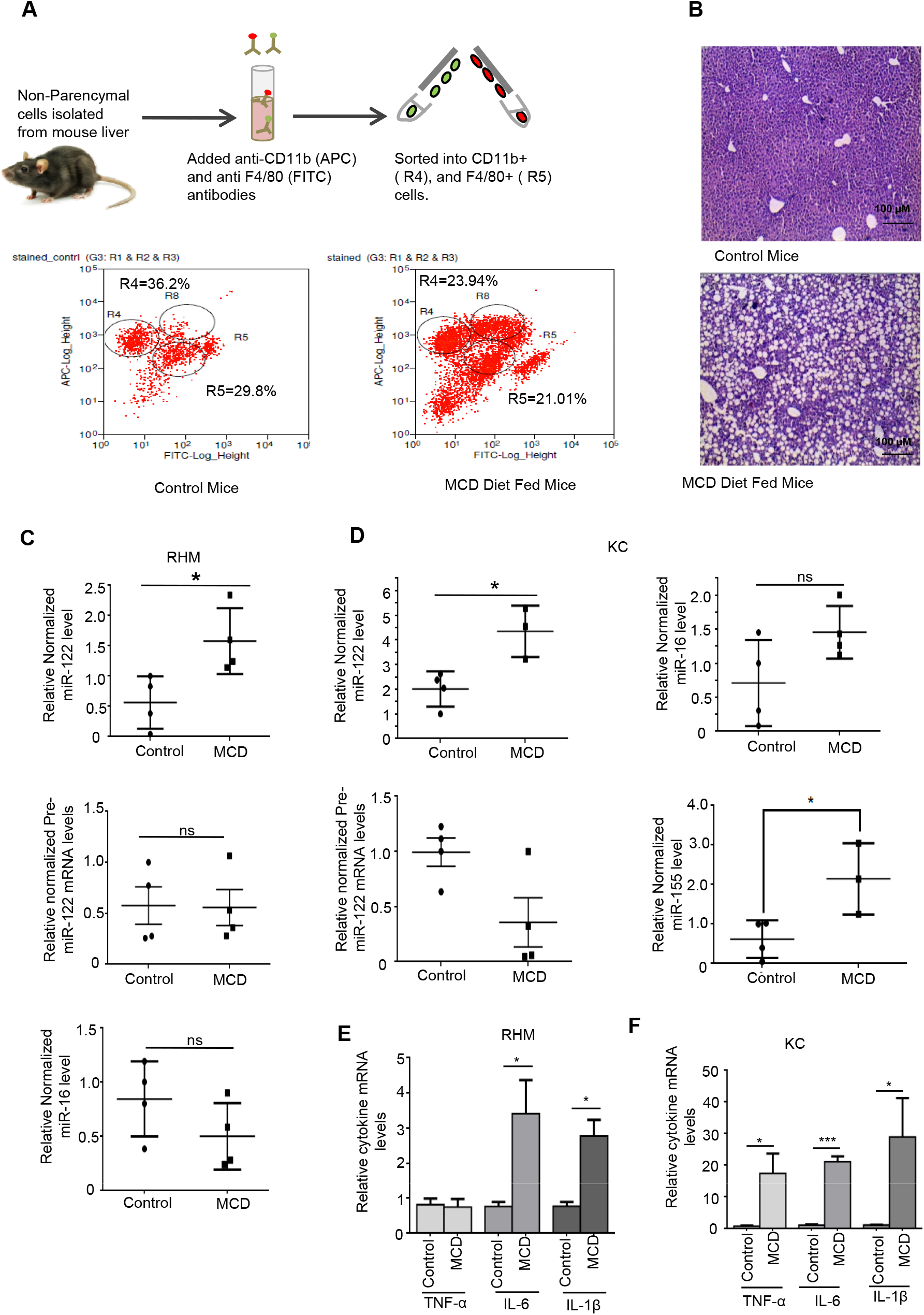
Transfer of miR-122 from hepatic cells to tissue macrophages in MCD diet fed mouse liver. **A** Isolation and characterization of Non-Parenchymal Cells (NPCs) from chow diet and MCD diet fed mouse liver. Schematic diagram showing isolation and FACS sorting of NPCs from chow diet fed (Control) vs MCD diet fed mice livers with CD11b (APC) and F4/80 (FITC) gating. **B** Haematoxylin and Eosin stained micrographs of control and MCD diet fed mice liver sections **C-D** Hepatic immune cells in MCD diet fed mice show significantly higher levels of internalized miR-122 with respect to control. Relative internalized miR-122 level in sorted RHM cells is shown (C, top panel). As a control, levels of pre-miR-122 (C, middle panel) and internalized miR-16 (C, bottom panel) have also been measured. Relative levels of internalized miR-122 in sorted KC cells is shown (D-top panel, left side). As a control, relative pre-miR-122 (D-bottom panel, left side) and internalized miR-16 (D-top panel, right side) and miR-155 levels (D-bottom panel, right side) have also been shown in the sorted KC cells. miR-155 induction is characteristic of inflammatory response. **E-F** Higher levels of internalized miR-122 correlates with increased expression of various pro-inflammatory cytokine mRNAs. Levels of pro-inflammatory cytokines (TNFα, IL-6 and IL1β) in sorted RHM cells (E) and KC cells (F) from control and MCD diet fed mice livers are shown. All mRNA and miRNA detections were done by RT-qPCR. RT-qPCR detection of cytokine mRNAs and pre-miR-122 was done using specific primers from 200 ng of cellular RNA. RT-qPCR detection of internalized miRNAs and miR-155 was done from 25ng of isolated cellular RNA. Normalization of miR-122, miR-155 and miR-16 were done against U6 snRNA. Normalization of precursor miR-122 and cytokine mRNAs were performed in respect to 18S rRNA levels. Data represents Mean ± SD. All images are representative of atleast 4 independent mice from each group. For statistical significance, minimum three independent experiments were considered in each case unless otherwise mentioned and error bars are represented as mean ± S.D. P-values were calculated by utilising Student’s t-test. ns: non-significant, *P < 0.05, **P < 0.01, ***P < 0.0001.

RT-qPCR analysis of RNA from isolated RHM (R4-F4/80^low^/CD11b^+^ set) showed higher levels of hepatic miR-122 in RHM of MCD diet fed mice livers as compared to those isolated from Chow diet fed mice (Figure 1C; upper panel). However levels of miR-16, another hepatic miRNA known to be present in circulation (Tan et al, 2014), remained unchanged (Figure 1C; lower panel). We also checked levels of precursor miR-122 levels in RHM to verify that miR-122 was not being endogenously induced to be expressed in the sorted RHM cells and was neither contributed by contamination from miR122 expressing hepatic cells (Figure 1C; middle panel). Equivalent results were obtained for KC (R5-F4/80^+^/CD11b^+^) cells sorted from Chow and MCD diet fed mice livers. Cells from MCD diet fed mice show significantly elevated levels of hepatic miR-122 (Figure 1D; upper left panel), whereas changes in miR-16 levels were found to be non-significant (Figure 1D; upper right panel). Sorted KC cells also do not induce transcription of pre-miR-122 as verified by decreased levels of precursor miR-122 in MCD diet fed mice as compared to Chow diet (Figure 1D, lower left panel). Interestingly, in the sorted KC cells there was an upregulation of miR-155, an inflammatory miRNA (Figure 1D, lower right panel).

Relative quantification of pro-inflammatory cytokine mRNAs was compared between control (Chow diet fed Mice) and MCD diet fed group. Internalization of miR-122 correlated with increased levels of IL-6 and IL-1β in RHM (F4/80^low^/CD11b^+^) from MCD diet fed mice livers (Figure 1E). Similar increases in TNFα, IL-6 and IL-1β levels were detected in KC (F4/80^+^/CD11b^+^ cells) cells (Figure 1F). Thus, both tissue resident KCs and recruited monocytes from circulation are in an inflammatory state in the obese liver where elevated presence of hepatic miR-122 was noted.

### Induction of inflammatory cytokines in tissue resident macrophages is caused by increased transfer of hepatic miR-122

How does the liver derived miR-122 increase in tissue resident macrophages? We wanted to investigate the role of lipid exposed liver cell derived extracellular vesicles (EVs) in the induction of inflammation of receiving macrophages. Huh7 is a human hepatoma cell known to express miR-122. These cells, when exposed to a cholesterol-lipid concentrate present in the culture medium for 4 hours, were found to export miR-122. EVs isolated from Huh7 cells (Basu and Bhattacharyya, 2014) were examined for their miRNA content (Figure 2B). As expected we detected increased levels of miR-122 there. Three other miRNAs, miR-16, miR-24 and miR-21 also showed increased levels in EVs isolated from cholesterol treated cells. miR-16 does show a relatively lower change in its level in released EVs after cholesterol exposure (Figure 2B). To test whether the EVs isolated from hepatic cells can increase the inflammatory response in recipient macrophage cells, we treated phorbol 12-myristate-13-acetate (PMA)-differentiated U937 macrophages with EVs isolated from cholesterol-lipid concentrate treated Huh7 cell. There, similar to what was noted in murine liver, addition of lipid induced hepatocyte-derived EVs elevated pro-inflammatory cytokine mRNA levels in human macrophage cells (Figure 2C).

**Figure 2.**
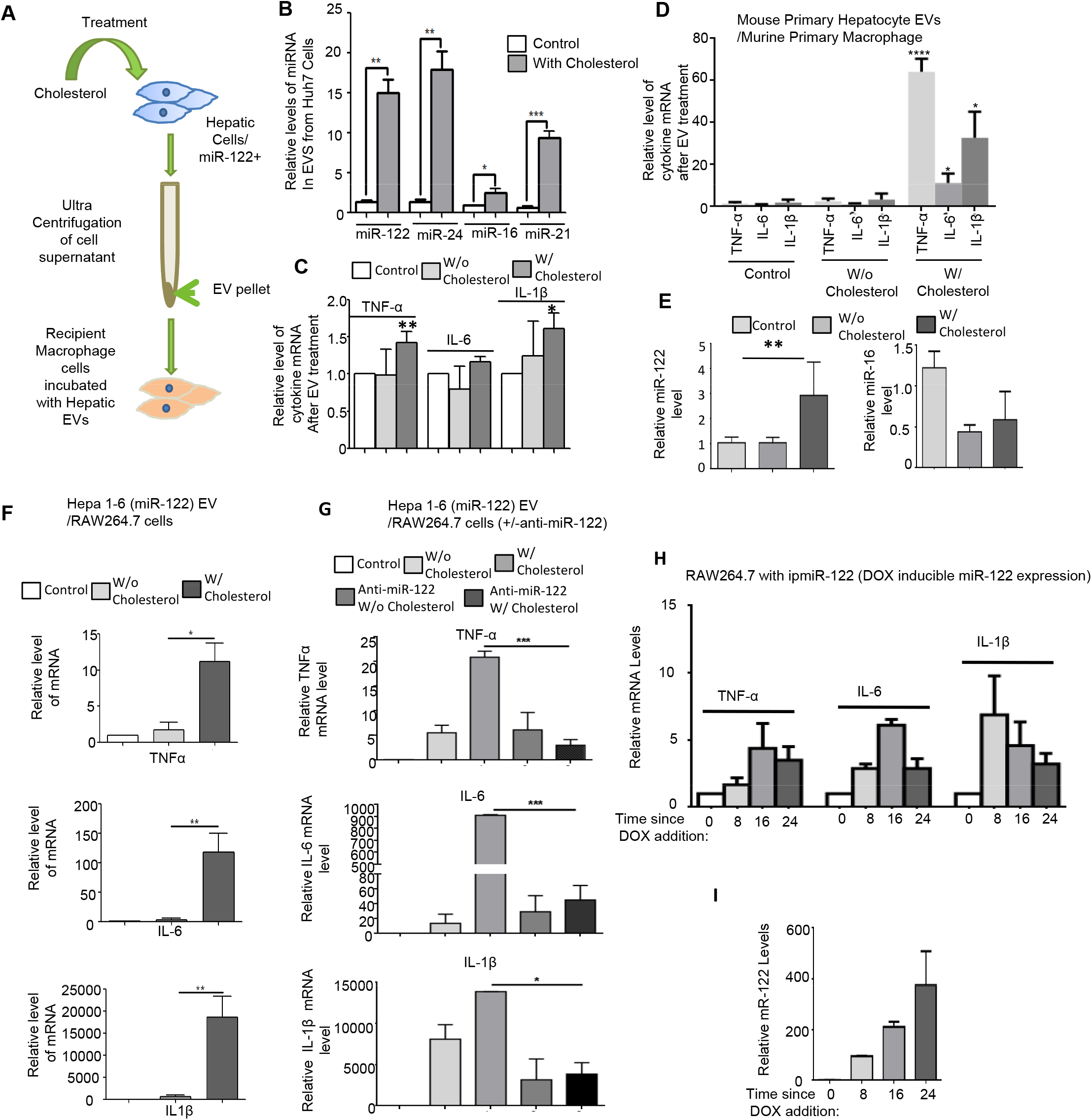
Extracellular Vesicle mediated transfer of miR-122 from lipid-exposed hepatic cells to tissue macrophages activates them to express pro-inflammatory cytokines. EV associated miR-122 from lipid treated hepatic cells when transferred to recipient macrophages induce higher levels of proinflammatory cytokine mRNAs. Exosomes were isolated from different types of hepatic cells treated with 5x cholesterol-lipid concentrate for 4 hours. For the ‘w/o cholesterol’ sets, equivalent volume of cholesterol–lipid concentrate was added to the cell supernatant <u>after</u> its collection from the cell culture plates. The ‘control’ set represents immune cells to which no EVs have been added. EV treatment was done for 16hours following which recipient cells were lysed and cellular RNA was isolated. **A** Schematic diagram showing experimental procedure described above. **B** Relative levels of exosomal miRNAs secreted by Huh7 cells upon exposure to cholesterol-lipid concentrate. **C** Relative levels of TNFα, IL-6 and IL-1β in PMA-differentiated U937 cells to which EVs isolated from cholesterol treated Huh7 cells were added. **D** Relative proinflammatory cytokine mRNA levels in murine primary macrophages incubated with EVs isolated from cholesterol treated primary hepatocytes. **E** Internalized miR-122 (left panel) and miR-16 (right panel) in primary macrophages of experiment described in (D). **F** Relative levels of TNFα mRNA (top panel), IL-6 mRNA (middle panel) and IL-1D mRNA (bottom panel) in recipient RAW 264.7 monocytes incubated with EVs from cholesterol treated Hepa1-6 cells expressing miR-122 (miR-122 expression plasmid pmiR-122 transfected). **G** Inhibition of internalized miR-122 in recipient macrophages leads to reversal of proinflammatory cytokine expression. EVs isolated from untreated (w/o cholesterol) and 5x cholesterol-lipid concentrate treated (w/cholesterol) Hepa1-6 cells (pmiR-122 transfected) were added to RAW 264.7 cells prior-transfected with anti-miR-122 oligos (Anti-miR-122 w/o cholesterol, anti-miR-122 w/ cholesterol). mRNA levels of TNF-α (top panel), IL-6 (middle panel) and IL-1β (bottom panel) were detected by RT-qPCR. **H-I** Presence of miR-122 in macrophage cells induces a pro-inflammatory phenotype. Induction of miR-122 expression in RAW264.7 leads to elevated levels of proinflammatory cytokine mRNAs. Tetracycline repressor expressing RAW264.7 cells were transfected with inducible miR-122 expressing plasmids. 400 ng/ml of Doxycycline was used to induce miR-122 expression. mRNA levels of TNF-α, IL-6 and IL-1β detected by RT-qPCR in cells harvested at indicated time points after Dox addition have been shown here (H). Relative miR-122 levels in RAW264.7 at various time points since DOX addition (I). All mRNA and miRNA detections were done by RT-qPCR. RT-qPCR detection of cellular miRNAs was done from 25 ng of cellular RNA. RT-qPCR detection of EV miRNAs was done from 100 ng of isolated EV RNA. Normalization of miR-122 and miR-16 were done with respect to U6 snRNA. Normalization of precursor miR-122 and cytokine mRNAs were done with respect to 18S rRNA. Data represents Mean ± SD. (B-I) N= 3 replicates. P-values were calculated by utilising Student’s t-test. ns: non-significant, *P < 0.05, **P < 0.01, ***P < 0.0001.

To determine the contribution of hepatic cell derived EVs in the induction of the inflammatory phenotype, mouse primary hepatocyte cells were first treated with cholesterol-lipid concentrate for 4 hours. EVs from the cell culture supernatant were then added to murine primary macrophages (Kupffer cells). Increased mRNA levels of the pro-inflammatory cytokines TNFα, IL-6 and IL-1β were observed in recipient Kupffer cells upon treatment with EVs derived from lipid treated primary hepatocytes (Figure 2D). RT-qPCR assays detected increased levels of internalized miR-122 in the recipient macrophage cells as compared to control (Figure 2E). However levels of internalized miR-16, another miRNA known to increase in circulation in high-lipid condition (Tan et al., 2014) remained unaffected (Figure 2E). To determine the miR-122 dependence of this phenomenon, we isolated EVs from cholesterol-lipid treated Hepa1-6 cells expressing miR-122 and added them to RAW 264.7 cells. Like in primary macrophage cells, RAW264.7 cells receiving the EVs from miR-122 transfected and cholesterol treated Hepa 1-6 cells also showed increased expression of pro-inflammatory cytokines (Figure 2F). In the subsequent experiment, to confirm the immunostimulatory role of miR-122 transferred from lipid challenged hepatic cells, receiving RAW264.7 cells were transfected with anti-miR-122 oligos. Anti-miR-122 transfected cells showed marked decrease in various proinflammatory cytokine mRNAs like TNF-α, IL-1β and IL-6 expression even after treatment with cholesterol-lipid treated Hepa1-6 EVs (Figure 2G). These results strongly suggest the pro-inflammatory nature of the miR-122 exported from lipid loaded hepatic cells. The extracellular miR-122, taken up by receiving macrophages, enhances the expression of proinflammatory cytokines and thereby ensures a more activated state of inflammation in the recipient cells.

To make a strong correlation of EV-derived miR-122 and inflammatory response, we expressed miR-122 from a doxycycline (DOX) inducible expression vector in naive RAW264.7 cells and checked cytokine mRNA levels over time when cells were incubated with DOX. The induction of miR-122 resulted in an accumulation of mature miR-122 over time upon induction with DOX. The increase in cellular miR-122 levels was found to correlate with the cytokine mRNAs’ expression increase (Figure 2 H and I). Interestingly after a threshold of expression, happening at 24h of miR-122 induction, effectiveness of miR-122 for cytokine induction gets impaired possibly due to jeopardized miRNP balance happening for other important miRNAs present there.

Do the EVs released by hepatic cells enter and affect the cytokine expression in tissue resident macrophages? We isolated EVs packed with miR-122 from the culture supernatant of HePa1-6 cells expressing miR-122. The EVs were injected into normal BALB/c mice (adult, 8-10 weeks) mice through tail-vein and hepatic non-parenchymal cells were isolated and analyzed for expression of different cytokines (Supplementary Figure S1A). Compared to PBS injected groups, mice injected with miR-122 containing EVs showed an increase in miR-122 levels in isolated hepatic non parenchymal cells, along with an elevation in inflammatory cytokines’ expression (Supplementary Figure S1B-C). The circulatory miR-122 level in miR-122 EV injected mice also showed an elevation with no corresponding change in serum miR-16 level (Supplementary Figure S1D). These data suggest miR-122 containing EV dependent inflammatory response in murine liver.

### Blocking of MMP2 affects miRNA cargo transfer between human cells

How do EVs move within the matrix of a tissue? In the liver, the extracellular matrix is rigid and composed of high levels of collagen. We expect, presence of specific proteases on the surface of EV may do the job to degrade the matrix and matrix metalloproteases (MMPs) are the best candidate for doing that. We were exploring the effect of different inhibitors and factors specific siRNAs against MMPs to score their effect on EV-mediated miRNA transfer in human cells in a microscopy based assay. In this assay the recipient HeLa cells were grown on coverslip in a 24 well cell culture format with the insert. The insert separated the upper compartment that was used for layering of Matrigel^R^. Isolated EVs from human hepatoma cell Huh7 transfected with Cy3-labeled miR-122 were used to study the transfer of Cy3 labeled miR-122 to recipient HeLa cells. The isolated EVs were added in the upper chamber which separated the EVs from target cells present in the lower chamber by a Matrigel^R^ layer. The transferred Cy3 labelled miRNAs were visualized by subjecting the recipient cell co-stained for tubulin to confocal microscopy imaging (Figure 3A). The inhibitors were applied in the upper chamber while SiRNA against specific proteins were used for co-transfection along with Cy3-miR-122 in donor Huh7 cells used for collecting EVs. The EVs used in the experiment were analyzed by Nanoparticle tracking analyzer (Figure 3B). We documented defective Cy3-miR-122 transfer happening in cells treated with ARP101 as an inhibitor of MMP2 (Jo et al, 2011). Similar results were also observed with EVs isolated from MMP2 deficient Huh7 donor cells (Figure 3C and E). Interestingly transfer of GFP-tagged CD63 positive EVs was also found to be impaired by ARP101 and siMMP2 when EVs were isolated from Huh7 cells expressing CD63-GFP (Figure 3D and F).

**Figure 3.**
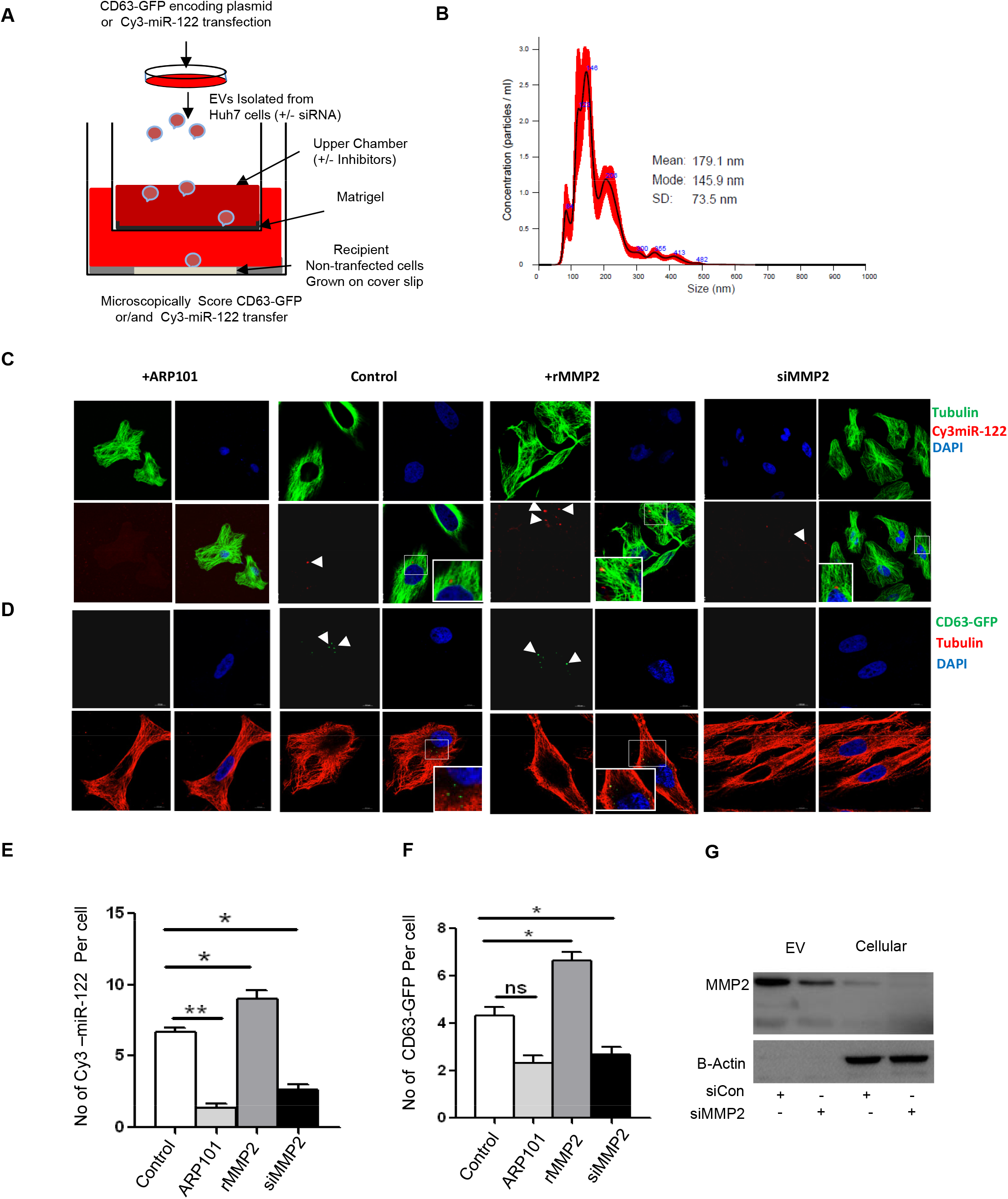
MMP2 dependent entry of EV in recipient cells. **A-C** Experimental scheme is shown in panel A. Donor Huh7 cells were transfected with cy3-labelled miR-122 oligonucleotides and siMMP2 before the EVs were isolated. The quantity and size distribution of the EVs were characterized by nano particle tracking analysis (B). Recipient HeLa cells were seeded on the gelatin coated coverslips at the lower chamber of the 24 well plates. Bottom of 0.4µM pore containing inserts were coated with diluted growth factor reduced matrigel (3mg/ml) and incubated in 37°C incubator for 30-45 min. After incubation, EVs were added into the inserts with the presence or absence of ARP101 or DMSO or rMMP2 and then it was kept for incubation for 24 hrs (A). Next day, after cell fixation, permeabilization and staining for β-tubulin cells were imaged (C). **D** Donor Huh7 cells were transfected with CD63GFP, siCon and siMMP2 and EVs were isolated. The number of CD63GFP transfer from donor cell exosome to recipient HeLa cell was also quantified by counting green dots of CD63 from three different sets, 5 fields/set, 5 cells/field. EVs from SiCon treated cells were used as control. ARP101 and rMMP2 were added along with EVs from CD63GFP expressing cells in respective condition. **E-G** The number of miR-122 transfer from donor cell exosome to recipient HeLa cell was also quantified by counting red dots of cy3-miR-122 from three different sets, 5 fields/set, 5 cells/field (E). The number of CD63GFP transfer from donor cell EVs to recipient HeLa cell was also quantified by counting green dots of CD63 from three different sets, 5 fields/set, 5 cells/field (F). The downregulation of MMP2 expression was observed when siMMP2 was transfected to donor cells by western blotting (G). For statistical significance, minimum three independent experiments were considered in each case unless otherwise mentioned and error bars are represented as mean ± S.E.M. P-values were calculated by utilising Student’s t-test. ns: non-significant, *P < 0.05, **P < 0.01, ***P < 0.0001. Scale bars in panel C and D are of 10 µm length.

### Functional transfer of miRNA from hepatic cells requires MMP2

What function does MMP2 have on EV-mediated cargo transfer between hepatic cells? To explore the exact role of MMP2 in miRNA transfer, we incubated miR-122 containing EVs derived either from naive or pmiR-122 transfected Huh7 cells with recipient HepG2 cells grown in Matrigel^R^. We documented a transfer of miRNA into HepG2 cells that otherwise don’t express miR-122. Further, we noticed blocking of this transfer of miRNA when incubated with ARP101, the MMP2 inhibitor. Treatment of donor Huh7 cells with anti-miR-122 blocks the transfer of functional miR-122 to recipient cells (Figure 4A). Similar level of miR-122 transfer from donor HeLa cells, ectopically expressing miR-122, to HepG2 has been documented and the MMP2 inhibitor blocked the transfer of miR-122 (Figure 4B). Do the transferred miRNA become functionally active in the recipient cells? We used miR-122 reporter mRNA to test the functional transfer of miR-122 in recipient cells. The transferred miR-122 could repress the RL-reporter having one perfect miR-122 sites in HepG2 recipient cells and a fold repression of the reporter has been scored against the RL reporter without miRNA binding sites (Figure 4C-D)(Basu & Bhattacharyya, 2014). Blocking of MMP2 stopped the miR-122 activity transfer to recipient HepG2 cells both from Huh7 and HeLa donor cells (Figure 4E and F). Interestingly, when EVs isolated from Huh7 cells expressing miR-122 were incubated with Huh7 cells we could see a drop in cellular miR-122 content. This may be explained by a reduced self-transfer of miR-122 to the donor cell that itself gets reduced in presence of MMP2 inhibitor ARP101 (Figure 4G). The results described here point out to an important role of MMP2 in transfer of functional cargo miRNA in mammalian cells and thus is an essential component of miRNA activity regulation in a tissue.

**Figure 4.**
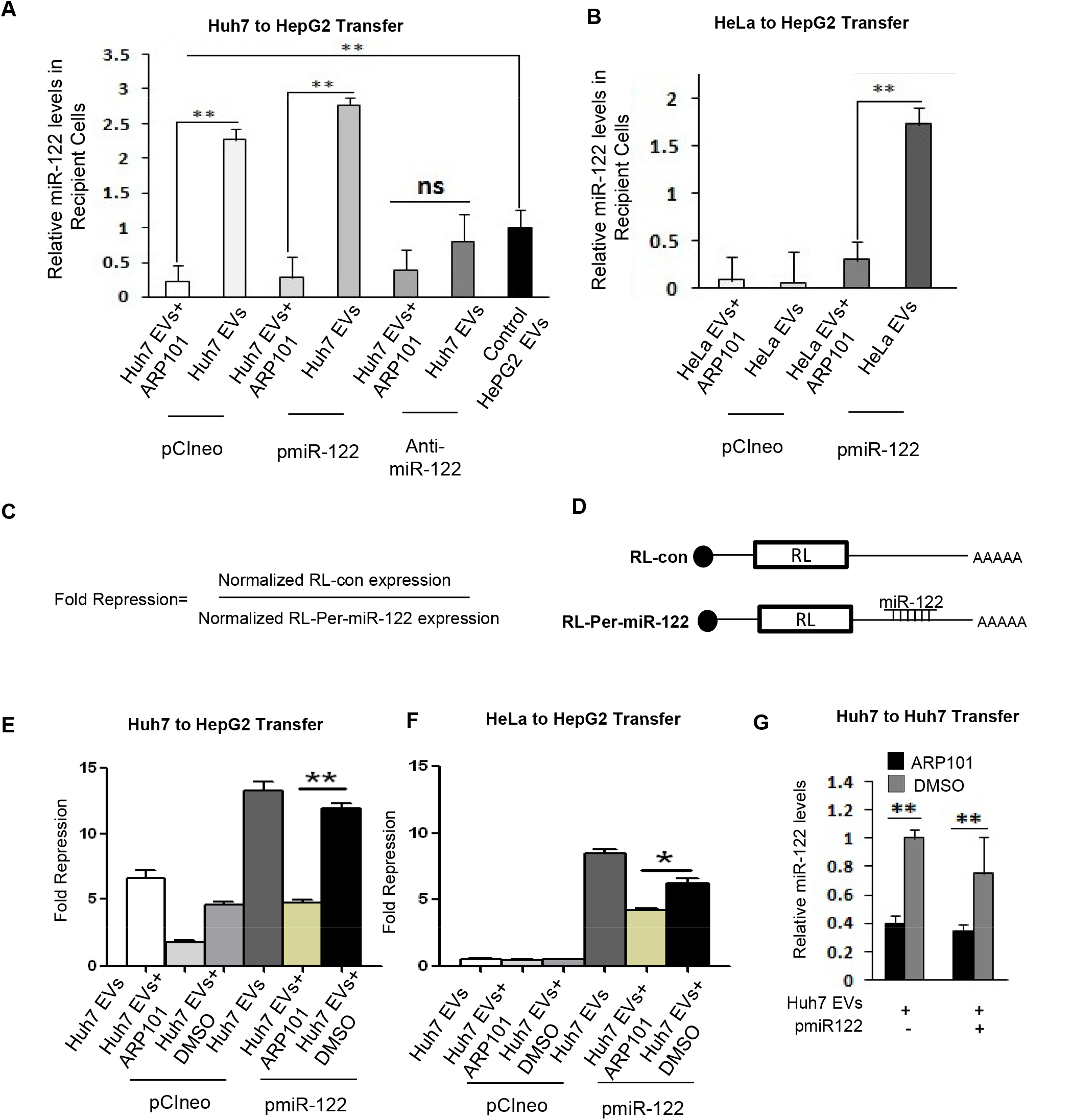
Functional transfer of miR-122 between hepatic cells requires MMP2 activity. **A-B** ARP101 inhibits the transfer of miR-122 containing EVs from donor cell to recipient cells. The transfer of EVs from hepatic cell lines were checked by using a specific MMP-2 inhibitor ARP101 (12.5µM/well). Huh7 (A) and HeLa (B) cells were chosen and transfected with miR-122 expressing pmiR-122 or anti-miR-122 oligoes and EVs were isolated. HepG2 cells were taken as recipient cell. To check the transfer, transwell inserts (0.4 µM, SPL) were taken and 100 µl 3mg/ml diluted matrigel were coated and solidified previously. In the lower chamber, HepG2 cells were seeded in complete DMEM media with 10% FCS. EVs were added in the medium in the inserts and incubated for 30 min in 37°C incubator. Serum free DMEM was added on to the inserts with ARP101. In control sets, same amount of DMSO (v/v) was added. This was then kept for 24 hrs in incubator. Next day, RNA was isolated by using Trizol™ RNA isolation method from HePG2 cells and cDNA was synthesized followed by real time PCR for miR-122 by using miR-122 primers. Normalization was done with respect to U6 snRNA. Data represents Mean ± SD. **C-F** Luciferase activity showed the repression of miR-122 reporter in recipient cells by transferred miR-122 from donor cells. The fold repression is the ratio of normalized expression levels obtained with RL-con and RL miR-122 reporter having one miR-122 perfect binding sites (C and D). In similar experiments described in A and B, the repressive activities in presence or absence of ARP101 were measured in HepG2 cells that don’t express miR-122 otherwise. Huh7, HepG2 and HeLa cells were overexpressed with miR-122 and exosomes were isolated and added to the matrigel coated 0.4micron inserts as previously. Recipient HepG2 cells were transfected with either RL-con or RL-per-miR-122 plasmids in parallel sets. To detect fold repression in HepG2, 10^6^ cells in a 10 cm^2^ well were transfected with 150 ng of each of the plasmids. Normalisation was done with a Firefly (FF) luciferase construct which was co-transfected along with the RL constructs (1µg for 1 × 10^6^ cells) after 24 h of transfection, cells were then spilted and seeded in the lower chamber of the transwell insert. ARP101 was added as described earlier. The whole set was incubated for 24 hrs again. Data represents Mean ± SD. **G** miR-122 expression was checked by ARP101 in donor cells. Huh7 cells were transfected with pmiR-122 and seeded in Matrigel with or without MMP2 inhibitor ARP101 in 24 well plate and incubated for 24 hrs. Cells were isolated from Matrigel and RNA was isolated followed by cDNA and Real time PCR for miR-122 and normalization of value obtained were done against U6 snRNA. Data represents Mean ± SD.

### MMP2 present on liver cell derived EVs and its association with EVs is dependent on MMP14

Presence of MMP2 with the exosomes isolated from human cells has been reported before where the MMP2 was found to promote the effect of exosomes on target cells (Tang et al, 2019). We have found presence of MMP2 in EVs isolated from liver and non-liver cells. We detected mature MMP2 in the EVs isolated from control Huh7 cells or cells expressing excess miR-122 (pmiR-122 transfected) and from Huh7 cells inactivated for miR-122 by anti-miR-122 treatment. In all cases we detected MMP2 in Gelatin zymography and by western blot (Figure 5A and B). We also detected MMP2 by western blot in EVs derived from non-hepatic cell HeLa (Figure 5B left panel). The mature form of the MMP2 was not detected in Huh7 and HeLa cells but mature MMP2 was predominantly found in the EVs. The EV associated MMP2 was found to be sensitive to Proteinase K treatment and no MMP2 was detected with EVs after proteinase K treatment (Figure 5C). These results suggest the presence of MMP2 on the outer side of the EVs. MMP14 is known to retain the MMP2 with cell and EV’s membrane (Han et al, 2015). To explore the retention mechanism of MMP2 with EVs, we inactivated MMP14 to check the presence of MMP2 associated with EVs. Depletion of MMP14 in Huh7 cells followed by isolation of EVs and western bloting for MMP2 showed a decrease in MMP2 levels in EVs isolated from MMP14 compromised Huh7 cells (Figure 5D). Our data suggest presence of MMP2 on the outer side of the EV membrane is dependent on MMP14 and the MMP2 is required for intercellular EV-transfer across the cell boundaries.

**Figure 5.**
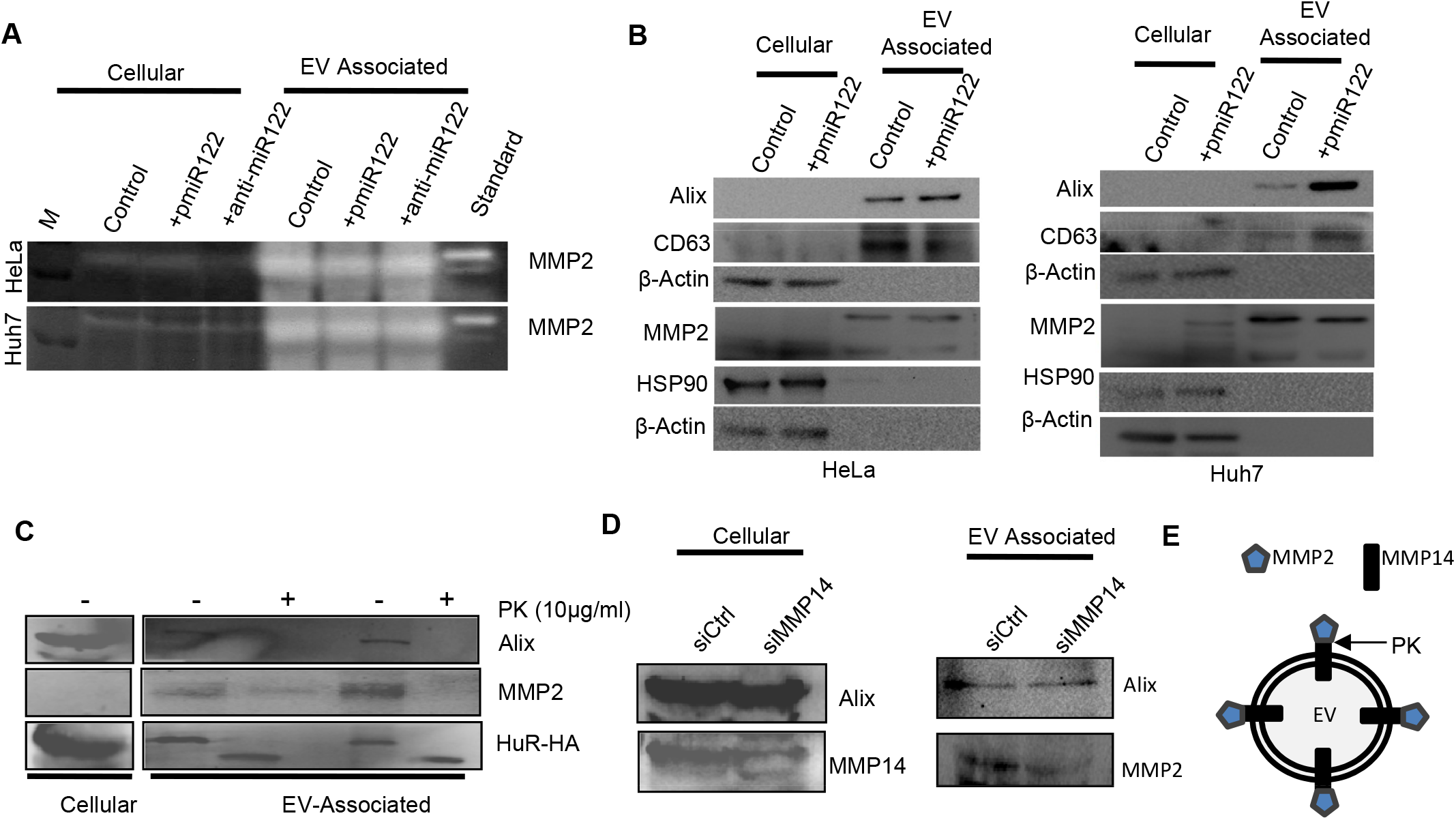
Presence of MMP2 on the Extracellular Vesicles (EVs) isolated from different hepatic and non-hepatic cells. A Gelatin Zymography reveals the presence of functional MMP2 in EVs isolated from hepatic cells. Extracellular vesicles or Exosomes were isolated from hepatic Huh7 cells transfected with or without miR-122 expressing plasmid pmiR-122 or pCIneo control plasmid. Anti-miR-122 treatment was also done in separate set cells. Cell lysates and EV extract (100ng protein each) were electrophoresed in 8% SDS-polyacrylamide gel containing 1 mg/ml gelatin under non-reducing conditions. B Presence of MMP2 in EVs isolated from hepatic and non hepatic cells (Huh7 & HeLa). Cells were transfected either with miR-122 expressing plasmid pmiR-122 or pCIneo vector. Extract of cells or isolated EVs (100 ng each) were then electrophoresed in 10% SDS-polyacrylamide gel. C Effect of Proteinase K (PK) treatment (20µg/ml) on EV associated Alix, MMP2 and HuR protein. Isolated EVS were treated with PK and after the incubation reaction was stopped and analyzed on 10% SDS-PAGE and western blotted. D-E Effect of MMP14 down regulation on MMP2 association with EVs. Huh7 cells were transfected with siRNAs specific to MMP14 or control siRNA and the cellular and EV associated levels of MMP14 and MMP2 were detected by western blot in respective sample (D). A suggested model of MMP14 mediated MMP2 recruitment of MMP2 to EVs (E)

### MMP2 facilitate EV movement across extracellular matrix and also enhance EV-entry in mammalian cells

Why MMP2 is required for the EV-cargo delivery? It is possible that MMP2 presence on EV enable them to migrate through the extracellular matrix by degrading the extracellular mesh made up of collagen and ensure a faster movement of the EV across the matrix. To test that, we used a collagen matrix and in a “EV-Movement” monitoring assay, we measured the EV movement across the matrix in presence and absence of ARP101 as MMP2 inhibitor or in presence of rMMP2. In control and rMMP2 presence we documented movement of EV front in collagen matrix measured by the rate of movement in unit time. While we documented no specific movement of EV front in presence of ARP101, confirming the importance of MMP2 in EV-movement across extracellular space (Supplementary Figure S2A-C). Does MMP2 also help in the internalization of EV? We did the EV-uptake experiment done in 2D cell culture without a Matrigel^R^ matrix. We noted the MMP2 dependent uptake of CD63-GFP positive EV entry in HeLa cells as the entry of CD63-GFP positive vesicles get retarded in presence of MMP2 inhibitor ARP101 (Supplementary Figure S2D-E). This data suggests a possible role of MMP2 in ensuring the movement of EVs across the matrix and its uptake in hepatic cells.

### Trafficking of miRNA containing EVs in mouse liver from hepatic cells to macrophage require MMP2

How does MMP2 affect the functional transfer of miRNA via EVs in a physiological context? miR-122 is a hepatocyte derived proinflammatory miRNA that when transferred to naive macrophage cells can activate them. To score the effect of MMP2 inhibition on transfer of hepatic EVs to resident macrophages, we adopted similar experimental set up as described in Figure 3A where the recipient macrophage cells were grown in the bottom chamber of a 12 well multi-well plate with insert and hepatocyte derived EVs were added on the upper chamber layered on top a Matrigel^R^ layer along with DMSO or ARP101 as MMP2 inhibitor. We scored the transfer of miR-122 to recipient Raw264.7 monocytes and measured the proinflammatory cytokine levels there (Figure 6A). We have documented upregulation of proinflammatory cytokines in cells where miR-122 were also getting transferred. Interestingly, blocking of miR-122 transfer by MMP2 inhibitor also affects the proinflammatory cytokine expression there (Figure 6A-B). To make a direct correlation between MMP2 and proinflammatory response in recipient RAW264.7 cells, we treated the EVs with rMMP2 and documented a positive influence of rMMP2 on proinflammatory response induced by hepatic EVs in RAW264.7 cells. (Figure 6B).

**Fig 6:**
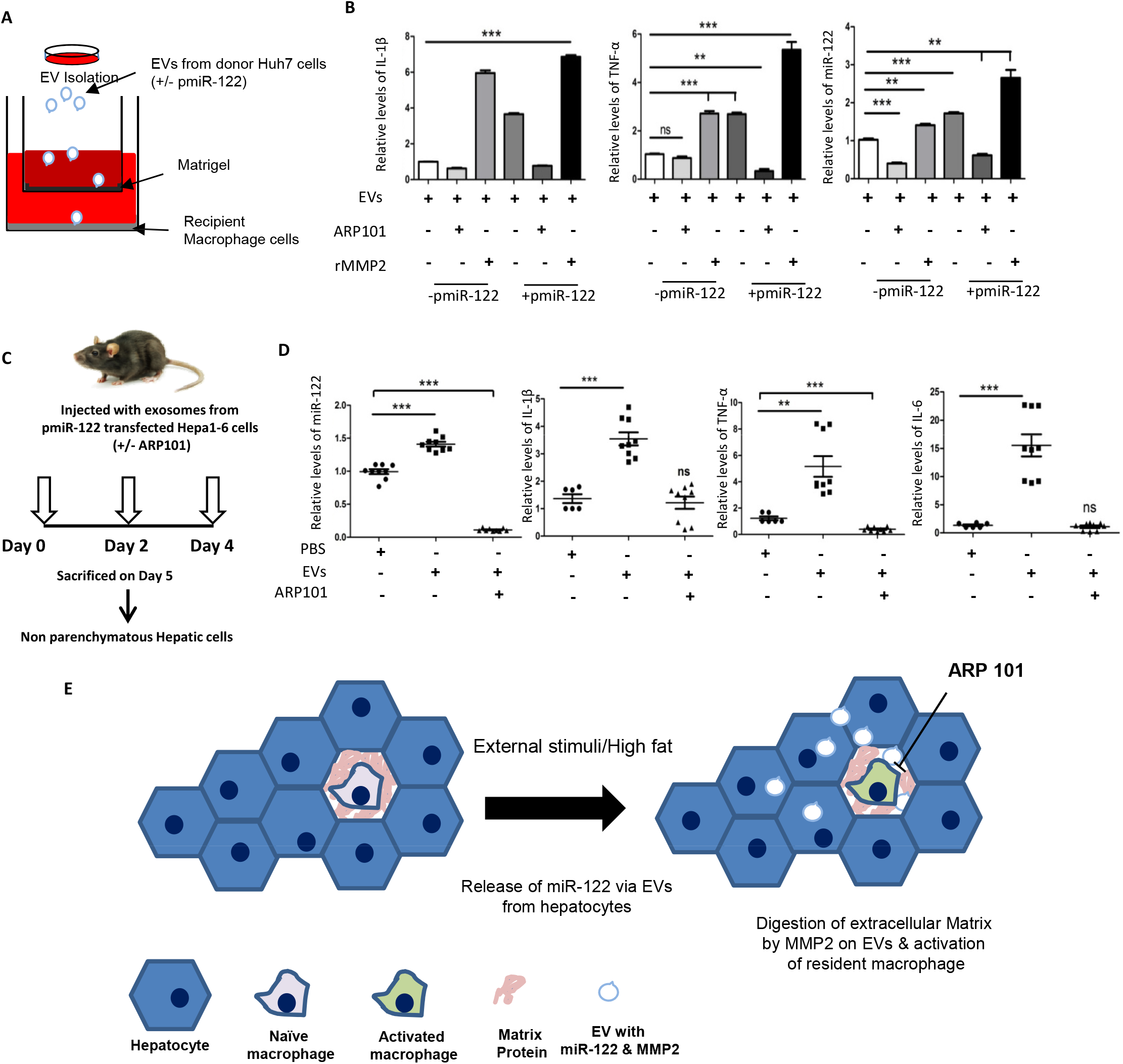
Inhibition of MMP2 affect transfer of pro-inflammatory signal across the hepatic cell boundary. MMP2 facilitates the miR-122 transfer to macrophage that leads to elevated levels of various pro-inflammatory cytokine mRNAs. Scheme of the experiment has been shown in panel A. Huh7 was transfected with control vector or pmiR-122 and EVs were isolated. Here RAW264.7 cells were taken as recipient cell. To check the transfer, transwell inserts (0.4 micron, SPL) were taken and 100 µl (3mg/ml) diluted matrigel were coated and solidified previously. In the lower chamber, RAW264.7 cells were seeded in complete RPMI media with 10% FCS. Then RAW264.7 cells were activated using LPS (1mg/ml) for 4 hrs. Exosomes were added on the inserts and incubated for 30 min in 37°C incubator. Serum free DMEM was added on to the inserts with ARP101 (12.5µM/well). In control sets, same amount of DMSO (v/v) was added. This was then kept for 24 hrs in incubator. Next day, inserts were removed and media from the lower chamber was discarded and RNA was isolated by using Trizol™ RNA isolation method. cDNA was synthesized followed by real time PCR for pro inflammatory (IL-1β and TNF-α) cytokines and miR-122 (B). Normalization was done with respect to 18S rRNA and U6 snRNA respectively. Data represents Mean ± SD. **C-D** Relative internalized miR-122 levels to non-parenchymatous hepatic cells in presence and absence of ARP101. The scheme of the experiment has been shown in panel C. EVs isolated from miR-122 expressing Hepa1-6 cells, were injected through the tail vein every alternate day for three days and mice sacrificed the next day after the third injection. Equal volume of PBS was administered as the control. Non-prenchymatous hepatic cells were isolated. Low miR-122 level was detected when MMP2 inhibitor was applied. Normalization was done with respect to U6 snRNA. Data represents Mean ± SD (D). Cytokine mRNA levels in isolated non-parenchymatous hepatic cells have been shown here. mRNA levels were detected by RT-qPCR. Normalization was done with respect to 18S. Data represents Mean ± SD (D). E A model of EV-mediated activation of resident macrophage in liver. The miR-122 conting EVS released by activated hepatocytes digest the matrix by associated MMP2 and get transferred to non-parenchymatous hepatic cells to activate them to express the pro-inflammatory cytokines-a process get inhibited by ARP101, the MMP2 inhibitor. For statistical significance, minimum three independent experiments were considered in each case unless otherwise mentioned and error bars are represented as mean ± S.E.M. P-values were calculated by utilising Student’s t-test. ns: non-significant, *P < 0.05, **P < 0.01, ***P < 0.0001.

To score the *in vivo* effect of the MMP2 inhibition in induction of inflammatory response by hepatic EVs with miR-122, we expressed miR-122 in Hepa1-6 cells and isolated the EVs. The EVS were injected through tail-vein injection in recipient mice alone or with MMP2 inhibitor ARP101. We documented reduced levels of pro-inflammatory cytokine expression and miR-122 transfer in non-parenchymal hepatic cell population in animals livers injected with ARP101 (Figure 6C-D). Our data suggests the importance of MMP2 for intra-tissue miRNA transfer in mouse liver.

## Discussion

In the work described above, we have identified how the matrix metalloproteinase MMP2 by facilitating the movement of EVs through the extracellular matrix affects the functional miRNA transfer in the hepatic context and plays a role in liver-specific miRNA-mediated inflammation. miRNA transfer between similar or different types of cell in a tissue causes miRNA homeostasis but how the miRNA expression levels are controlled by the factors that affect the transfer process per se are unknown.

Extracellular vesicles (EVs) help in cell to cell communication and, like hormones, can have an autocrine, paracrine or endocrine effect (Becker et al, 2016). miRNAs packaged in EVs are speculated to fine tune the gene expression profile in neighbouring cells and tissues and thus, could facilitate metabolic and functional homeostasis in respective tissues (Chevillet et al, 2014; Lotvall & Valadi, 2007). In the current work, miR-122 enriched EVs secreted from lipid loaded Huh7 cells have been shown to elicit a pro-inflammatory response in recipient macrophages. Is there a reciprocal effect of secreted pro-inflammatory cytokines on Huh7 cells? The released cytokines may bind to receptors on Huh7 cells and affect the transcription flux of HuR other factors that mediate the extracellular export pathway (Mukherjee et al, 2016). Possibility of the existence of such a feedback loop could help to curb the pro-inflammatory response in macrophages and could also, fine tune the miRNA profile in Huh7 cells and, thus could help in the attainment of tissue level homeostasis in gene expression. Deciphering the mechanism of such a feedback loop can be looked into in future studies. The propagation of inflammatory miR-122 as part of EVs in the blood of high fat diet fed mice raises the possibility of the miRNA being taken up by far located tissue macrophage in other organs where it may play a role in ectopic activation of the immune cells across the tissue boundary and thus, contribute to a chronic systemic inflammatory response in mice exposed to high-fat diet.

miRNA transfer can be controlled at three different steps; the packing and export of the miRNAs via EVs, the movement of EVs through the extracellular matrix and internalization and release of content of EVs in the recipient cells. Exploration of all three steps is largely limited in identification of the few factors individually controlling these steps. HuR is one such protein that has been identified as the facilitator of export of specific subset of miRNAs including miR-122 from hepatic cells to ensure its export in stress conditions (Mukherjee et al, 2016). This protects the hepatic cells from stress. The effect of stress on MMP2 has been known and it also seems to have limited effect on MMP2 present on the EVs released by the hepatic cells under stress condition (data not shown). From the data described here, MMP2 expression also seems to remain unaffected in Huh7 cells at protein level after expression of excess miR-122. miR-122 after getting into hepatic cells also decreases factors preventing its expression such as hepatic Insulin like growth factor 1 to ensure robust miR-122 expression (Basu & Bhattacharyya, 2014).

miRNAs, as epigenetic signals should exchange between neighbouring cells in a functional form and to ensure that they are transferred as single stranded form and getting incorporated into recipient cell Ago2 protein (unpublished data) and plays an important role in controlling the expression of target mRNA in both donor and recipient cells. To ensure the effective transfer of miRNAs, the movement of EVs through the matrix is an important factor and several biochemical and physical properties of the matrix should have contributed to controlling the speed of the movement. The collagen network and extracellular proteases must work in a reciprocal manner to ensure the movement. The concentration and maturation of MMPs should be the key aspects that should determine the EVs movement. In tumour microenvironment, the multidimensional movement of cancer cell derived EVs contributes in the establishment of cancer cell niche by affecting the gene expression of non-cancerous and immune cells present in the tumour. Variable expression of MMPs by cancer cells are known to be essential for niche creation. We propose the MMP-mediated facilitation of the EV’s movement may thus contribute to cancer progression by ensuring a rapid movement of EVs in 3D. Blocking of EV-transport by targeting MMPs would thus be an effective way of controlling miRNA transport in mammalian cells and may be an useful tool to curtail cancer tumour growth.

## Experimental Procedures

### Cell Culture, Reagents and Antibodies

Human HCC cell lines (Huh7, HepG2) and Human HeLa were cultured in Dulbecco’s modified Eagle’s medium (DMEM; Gibco-BRL) supplemented with 10% fetal bovine serum (FBS; GIBCO-BRL) and Penicillin Streptomycin (1X) antibiotics (GIBCO). RAW264.7 were cultured in RPMI (Gibco-BRL) supplemented with 10% fetal bovine serum (FBS; GIBCO-BRL) and Penicillin Streptomycin (1X) antibiotics (GIBCO).

MMP2 inhibitor, ARP101 was purchased from Sigma-Aldrich; recombinant MMP2 was purchased from Calbiochem. DMSO was obtained from Fisher Scientific.

Human Alix (Santa Cruz), CD63 (BD Pharmingen), MMP2 (Cell Signaling Technologies), Actin (Sigma-Aldrich) and secondary mouse and rabbit antibodies (Invitrogen) were purchased.

### Plasmid constructs, Cell transfections and Luciferase assay

The RL reporters (Renilla luciferase) were previously described (Pillai et al., 2005). siControl and siMMP2 were used for transfection at 100 pmoles per well of a confluent six-well plate. miR-122 and anti-miR-122 were purchased from Ambion and was used at 100 pmoles to transfect cells per well of a six-well plate. Cells were differentially transfected microscopy using CD63-GFP, Tubulin-GFP and cy3-miR122 plasmids. For transfections 1µg of the plasmids was used for transfecting 10^6^ cells in a 10cm^2^ well. All transfections were performed using Lipofectamine 2000 (Invitrogen) following manufacturer’s instructions. For luciferase assays 10^6^ cells in a 10 cm^2^ well were transfected with either RL-con or RL-per-miR-122 plasmids in parallel sets. To detect fold repression in HepG2, 10^6^ cells in a 10 cm^2^ well were transfected with 150 ng of each of the plasmids. Normalisation was done with a Firefly (FF) luciferase construct which was co-transfected along with the RL constructs (1µg for 1 × 10^6^ cells) after 24 h of transfection, cells were splited After that cells were lysed with 1 X Passive Lysis Buffer (Promega). Renilla (RL) and Firefly (FL) activities were measured using a Dual-Luciferase Assay Kit (Promega) following the suppliers protocol on a VICTOR X3 Plate Reader with injectors (Perkin Elmer). Mean Fold Repression was calculated by dividing the FF normalized RL-Con value with that of FF normalized RL-per-miR-122 value. Relative fold repression was calculated by taking the control mean fold repression as 1. All luciferase assays used in this study have been done in triplicate. All experiments were performed minimum three times before the SD values were calculated.

### Cholesterol and Palmitic Acid treatment

MβCD conjugated cholesterol conjugate obtained from GIBCO (#12531-018) was added from a 250X stock to Huh7 cells in culture at a final concentration of 5X for a period of 4 hours. Cholesterol treatments were done to Huh7 cells in fresh growth media at 70-80% confluency. Unless otherwise mentioned, cholesterol treatment was done at a final concentration of 5x for 4 hours.

### Animal experiments

All animal experiments were approved by the Institutional Animal Ethics Committee (approved by CPCSEA, Ministry of Environment & Forest, and Government of India). 8-10 weeks old male (20-24g) C57BL/6 mice were housed under controlled conditions (temperature 23 ± 2c, 12 hour/12-hour light/dark cycle) in individually ventilated cages. Mice were randomly divided into two groups and fed either standard chow diet or methionine and choline deficient diet MCD (MP Biomedicals; #0296043910) up to four weeks.

C57BL/6 male mice of 8-10 weeks age were divided in two groups for normal chow and high fat diet (HFD) containing 45% fat and 5.81 kcal/gm diet energy content (MP Biomedicals; # 960192). Animals were fed with HFD for 4 weeks.

For isolation of RNA from tissues, TRIzol® (Invitrogen) reagent was used. For analysis of EV-associated RNA, serum fraction of blood was used. Relative levels of miRNA and mRNA in serum and tissues were quantified by qRT–PCR.

For histological analysis, tissues were fixed in 10% formaldehyde in PBS, embedded in paraffin, sectioned at 10 μm, and stained with hematoxylin and eosin (H&E) following standard staining protocol.

For exosome injection experiments, Hepa1-6 cells were transfected with miR-122 expressing plasmids (6μg/ 6 x 10^6^ cells). Approximately 1 x 10^9^ exosomes (measured by Nanoparticle Tracking Analysis-Nanosight Malvern U.K.) isolated from transfected Hepa1-6 cells (∼ 1x 10^7^ cells) were suspended in 1x PBS (passed through 0.22μm filter units in 100 µl) and injected into the tail vein of BALB/c mice (adult, 8-10 weeks). Control mice were injected with equal volume of 1x PBS (passed through 0.22μm filter units). Experiment was performed with 5 mice in each group (control injected and exosome injected). Injections were repeated every alternate day for three days and mice sacrificed the next day after the third injection. Animals were anaesthetized and livers were slowly perfused initially with the HBSS and then with 0.05% collagenase buffer via the portal vein. Livers were then excised, minced and filtered through a 70 μm cell strainer. The resultant single cell suspension was centrifuged at 50Xg for 5 minutes to precipitate and remove the hepatocytes. The supernatant was then collected and centrifuged at 250g for 15 minutes. The pellet was resuspended in RBC lysis buffer and kept in ice for 10 minutes. The resultant cell suspension was again centrifuged at 250xg for 15 minutes and pellet obtained was lysed with Trizol reagent. For detection of serum miRNA levels, serum fraction of blood was used. Blood samples were collected by cardiac puncture and allowed to clot. Serum was separated by centrifugation and frozen at −80°C.

### Liver macrophage sorting and analysis by flow cytometry (FACS)

Animals were anaesthetized and livers were slowly perfused initially with the HBSS and then with 0.05% collagenase buffer via the portal vein. Livers were then excised, minced and filtered through a 70μm cell strainer. The resultant single cell suspension was centrifuged at 250 g for 5 minutes to obtain non parenchymal cells and on the basis of surface staining with anti-F4/80 (FITC), anti-CD11b (APC) (eBioscience) antibodies, three types of liver macrophages were sorted by FACS (Beckman Coulter).CD11b+, F4/80+ and Cd11b+F4/80+ cells were sorted. Unstained cells were used for setting compensation and gates.

### Primary Hepatocyte Isolation

Animals were obtained from the animal house of the institute and all experiments were performed according to the guidelines set by Institutional Animal Ethics committee following the Govt. of India regulations. Mouse primary hepatocytes were isolated using the hepatocyte product line from Gibco Invitrogen Corporation. Adult BALB/c mice (4-6 weeks) were anaesthetized and the portal vein was cannulated using a 25G butterfly cannula and an incision was made in the inferior vena cava. The liver was perfused with 350 mL of warm (37°C) Liver Perfusion Medium (Cat. No. 17701) at a rate of 35 mL/minute with the perfusate exiting through the severed vena cava. This was followed by a Collagenase-Dispase digestion with Liver Digest Medium (Cat no. 17703) at a rate of 35 mL/minute. The liver was then aseptically transferred to the tissue culture hood on ice in Hepatocyte Wash Medium (cat no. 17704). Using blunt forceps the digested liver was torn open to release the hepatocytes. Cell clumps were dissociated by gently pipetting the solution up and down using a 25ml pipette. The solution was then filtered through 100 μM nylon caps atop 50 ml conical tubes. The cell suspension was then centrifuged at 50 x g for 3 min. The pellet was gently resuspended in 10 ml of Wash Medium using 25 ml pipette and the centrifugation repeated.

Cells were finally resuspended in Hepatocyte Wash Medium with 10% FCS and plated at 1×10^7^ cells /ml. Cells were plated in tissue culture treated collagen (Gibco Cat. No. A10483-01) coated plates at 12.5 µg/cm^2^. Unattached cells are poured off 4h after plating and medium was replaced with Hepatozyme-SFM (Cat no. 17705) with glutamine and 1% Pen/Strep. Cholesterol and BSA-Palmitate was added the next day in Hepatozyme-SFM.

For Kuppfer cell isolation, supernatant obtained from the first centrifugation (at 50x g for 3 minutes) after filtering through the 100 μm cell strainer was further centrifuged at 250xg for 5 min. The cell pellet was washed, resuspended in DMEM with 10% FBS, seeded onto tissue culture plates, and allowed to adhere for 16 h. Nonadherent cells were removed by several washes with PBS; >80% adherent cells were found to be positive for F4/80, a well-established KC marker.

### Exosome Isolation

For exosome isolation cells were grown in media made from exosome depleted FCS which were prepared by ultracentrifugation of the FCS used at 110,000×g for 5 h. The supernatant CM from one 90 cm^2^ plates, having 1×10^6^ donor cells (Huh7) was taken. The CMs were centrifuged first at 300×g for 10 min, then at 2000×g for 15 min followed by centrifugation at 10,000×g for 30 min. All centrifugations were done at 4^°^C. The CM was then filtered through a 0.22 m filter unit. This was then centrifuged at 100,000×g for 90 min at 4^°^C. After centrifugation, the supernatant was discarded. The pellet was resuspended in media and added back to recipient cells (HepG2) in a 24-well format such that 1 × 10^5^ recipient cells received the exosomes from 1 × 10^6^ donor cells. For CM based assays the same ratio was followed with CM from 10^6^ cells being added to 2 × 10^5^ cells. For the isolation of miR-122, anti-miR-122, Huh7, HeLa and HepG2 carrying exosomes, 1 × 10^6^ cells were transfected and 24 h after transfection, the cells were reseeded onto a 90 cm^2^ plate. Cells were grown for 48–72 h and exosomes isolated from the CM of these cells. For the exosomal protein isolation experiment, the CMs were centrifuged first at 300 x g for 10 mins, then at 2000 x g for 15 min followed by centrifugation at 10,000 x g for 30 mins. All centrifugations were done at 4°C. The CM was then filtered through a 0.22 µm filter unit and was loaded on a sucrose cushion (1 M sucrose and 10 mM Tris–HCl pH 7.5). This was ultracentrifuged at 120,000 x g for 90 mins at 4°C. The medium above the sucrose cushion was discarded leaving behind a narrow layer of medium with the exosomes at the interface. 1XPBS was added and the separated exosomes were washed at 4°C for 90 mins at 100,000 x g. The pellet was resuspended in 200µl of 1X SDS Buffer, which was used for western analysis for exosome markers.

### Gelatin Zymography and invitro EV Movement assay

Human HCC cells (Huh7 and HepG2) and a non hepatic cell line HeLa were transfected with miR122 expressing plasmid PJC148 and anti-miR122 oligonucleotides, media was changed after 6 hours and incubated for o/n at 37°C humidified incubator. Cells were splited to 90mm culture dishes containing DMEM with 2% Exo depleted FCS and kept for 24 hrs at incubator. Exosomes were isolated and EDTA free protease inhibitor cocktail containing lysis buffer was added to isolated exosomes and 8% gelatin containing SDS-PAGE gel was run at 60v for 3 hrs. After the completion of the run gels were washed for 1hr in 2.5x Triton X and then gels were incubated in calcium assay buffer (40 mM Tris-HCl, pH 7.4, 0.2 M NaCl, 10 mM CaCl_2_) for o/n at 37°C incubator. After incubation in buffer, gels were stained with 0.1% coomasie blue and then destaining was followed. The zones of gelatinolytic activities appeared as negative staining. Images of zymographic bands were performed using an UVP BioImager 600 system equipped with VisionWorks Life Science software (UVP) V6.80.

For the diffusion rate measurement assay, Huh7 released EVs were pretreated with rMMP2 or ARP101. EVS were isolated from culture supernatant of Huh7 cells (1×10^7^ Cells) and were then spotted on a well formed in the middle of a 35mm Petri dish layered with 2mm thick gelatine matrix. After 30 minutes the reaction was stopped by placing the gel for staining with Brilliant Blue to visualize the diffusion front of the EVs through the matrix. The representative plate photographs were taken and distance travelled by the MMP2 containing EVs were measured from the center by visualizing and measuring the distance of the front visible in individual cases.

### Western blotting

Cells were lysed in 1X Passive Lysis Buffer (PLB) (Promega) and quantified using Bradford reagent (Thermo Scientific). Cell number equivalent amount of the sample is then diluted in 5X Sample Loading Buffer (312.5 mM Tris-HCl, pH 6.8, 10% SDS, 50% glycerol, 250 mM DTT, 0.5% bromophenol blue) and heated for 10 mins at 95°C. Usually 1/5^th^ of the total amount from a 4cm^2^ well was loaded for both the control and experimental samples. Following SDS-polyacrylamide gel electrophoresis of the extracts, proteins were transferred to PVDF nylon membranes. Membranes were blocked in TBS (Tris-buffered saline) containing 0.1% Tween-20 and 3% BSA. Primary antibodies were added in 3% BSA for a minimum 16 hr at 4°C. Following overnight incubation with antibody, the membranes were washed at room temperature thrice for 5 min with TBS containing 0.1% Tween-20. Washed membranes were incubated at room temperature for 1 hr with secondary antibodies conjugated with horseradish peroxidase (1:8000 dilutions). Excess antibodies were washed three times with TBS-Tween-20 at room temperature. Antigen-antibody complexes were detected with West Pico Chemi luminescent substrate using standard manufactures protocol (Perkin Elmer). Imaging of all western blots was performed using an UVP BioImager 600 system equipped with VisionWorks Life Science software (UVP) V6.80.

### Fluoroscence Microscopy

Gelatin coated cover slips are added onto a 24 well plate. Recipient HeLa cells were seeded onto those cover slips such that they become 40% confluent after 24 hours. Exosomes from donor cells were added to 24-well inserts coated with 3mg/ml Matrigel and incubated for 24-48hrs. The recipient cells were washed with 1X PBS and fixed using 4% paraformaldehyde in 1X PBS for 30 mins in the dark at room temperature. Cover slips were then washed thrice with 1X PBS. Primary antibody incubation was done in 1XPBS with 1% BSA at 4°C overnight in a humid chamber. The anti-Tubulin antibody was used at a dilution of 1:100. Secondary antibody incubation was done in 1XPBS with 1% BSA for 1h at room temperature. Secondary anti-mouse antibodies labeled either with Alexa Fluor® 488 secondary antibodies (green) or Alexa Fluor® 568 (Red) (Invitrogen) were used at 1:500 dilutions. The cells were subsequently washed thrice with 1X PBS. Coverslips were then mounted with Vectashield containing DAPI and observed under a fluorescence microscope. Images were captured with a Zeiss LSM800 microscope. All post capture analysis and processing were done using Imaris 7 (BitPlane) software.

### Post Capture Image analysis

All western blots were processed with Adobe Photoshop CS4 for all linear adjustments and cropping. All images captured on Zeiss LSM800 microscope were analyzed and processed with Imaris 7 (Bitplane) software.

### Quantitative estimation of mRNA and miRNA levels

RNA was extracted by using the TRIzol reagent according to the manufacturer’s protocol (Invitrogen). Real time analyses by two-step RT-qPCR was performed for quantification of miRNA and mRNA levels. All mRNA RT-qPCRs were performed on a 7500 REAL TIME PCR SYSTEM (Applied Biosystems). mRNA real time quantification was generally performed in a two step format using Eurogentec Reverse Transcriptase Core Kit and MESA GREEN qPCR Master Mix Plus for SYBR Assay with Low Rox kit from Eurogentec following the suppliers’ protocols. Reactions were performed with 50ng of cellular RNA. The RT reaction condition was 25°C, 10 min; 48°C, 30min; 95°C, 5 min. The PCR condition was 95°C, 5 min; 95°C, 15 sec; 60°C, 1 min; for 40 cycles. The comparative C_t_ method which typically included normalization by 18S rRNA levels for each sample was used for relative quantification. Details of mRNA gene specific primers are given below.

Quantification of miRNA levels was done using Applied BiosystemTaqMan® chemistry based miRNA assay system. All mRNA RT-qPCRs were performed on Biorad CFX96 Real Time System. Assays were performed with 25 ng of cellular RNA, using specific primers for human miR-122, miR-16 and miR-21 (assay ID 000445, 000391, 000397 respectively). U6 snRNA (assay ID 001973) was used as an endogenous control. One third of the reverse transcription mix was subjected to PCR amplification with TaqMan® Universal PCR Master Mix No AmpErase (Applied Biosystems) and the respective TaqMan® reagents for target miRNA. The RT reaction condition was: 16°C, 30 min; 42°C, 30 min; 85°C, 5 min; 4°C, D. The PCR condition was: 95°C, 5 min; 95°C, 15 sec; 60°C, 1 min; for 40 cycles. For detection of miRNAs in exosomal fractions, 100ng of RNA was used for reverse transcription.

Samples were analyzed in triplicates. The concentrations of intra cellular miRNAs and mRNAs were calculated based on their normalized *C*_*t*_values. The ΔΔCt method for relative quantitation (RQ) of gene expression was used and relative quantification was done using the equation 2^−ΔΔCt^ (as per ‘Guide to Performing Relative Quantitation of Gene Expression Using Real-Time Quantitative PCR’ obtained from the Applied Biosystems website (http://www3.appliedbiosystems.com/cms/groups/mcb_support/documents/generaldocuments/cms_042380.pdf).

### mRNA RT-qPCR Primer Sequences

Mouse TNFα: Forward: 5’-GTCTCAGCCTCTTCTCATTCC-3’; Mouse TNFα: Reverse: 5’-TCCACTTGGTGGTTTGCTACG-3’; Mouse IL6: Forward: 5’-AGGATACCACTCCCAACAGA-3’; Mouse IL6: Reverse: 5’-GTACTCCAGAAGACCAGAGGA-3’; Mouse IL1β: Forward: 5’-GACCTTCCAGGATGAGGACAT-3’; Mouse IL1β: Reverse: 5’-CCTTGTACAAAGCTCATGGAG-3’; 18S rRNA: Forward: 5’-TGACTCTAGATAACCTCGGG-3’; 18s rRNA: Reverse: 5’-GACTCATTCCAATTACAGGG-3’.

### Statistical Analysis

All graphs and statistical analyses were generated in Graph-Pad Prism 5.00 (GraphPad, San Diego, CA, USA). Nonparametric unpaired t-test and paired t test were used for analysis, and P values were determined. Error bars indicate mean ±SEM.

## Acknowledgements

We thank Witold Filipowicz and Gunter Meister for different constructs used in this study. We thank the Funding body, Dept. of Science and Technology (DST), Govt. of India and Council for Scientific and Industrial Research (CSIR), and University Grant Commision (UGC) for the fellowship to DB, SB, S.C and MA. SB and D.D. also acknowledges the support from Department of Biotechnology, Govt of India and Dept. of Science and Technology (DST), Govt. of India for their fellowship support. SNB was supported by SwarnaJayanti Fellowship and a High Risk High Reward Grant from DST.

## Author Contributions

S.N.B. conceived the idea, designed the experiments, analyzed the data and wrote the manuscript. S.B., A.D., S.S and P.C. have contributed in design and planning the experiments. A.D. S.B.,S.C., D.D., and M.A. performed the experiments. S.B. and D.B also wrote the manuscript with S.N.B. and analyzed the data.

---

**Supplementary Figure S1.**
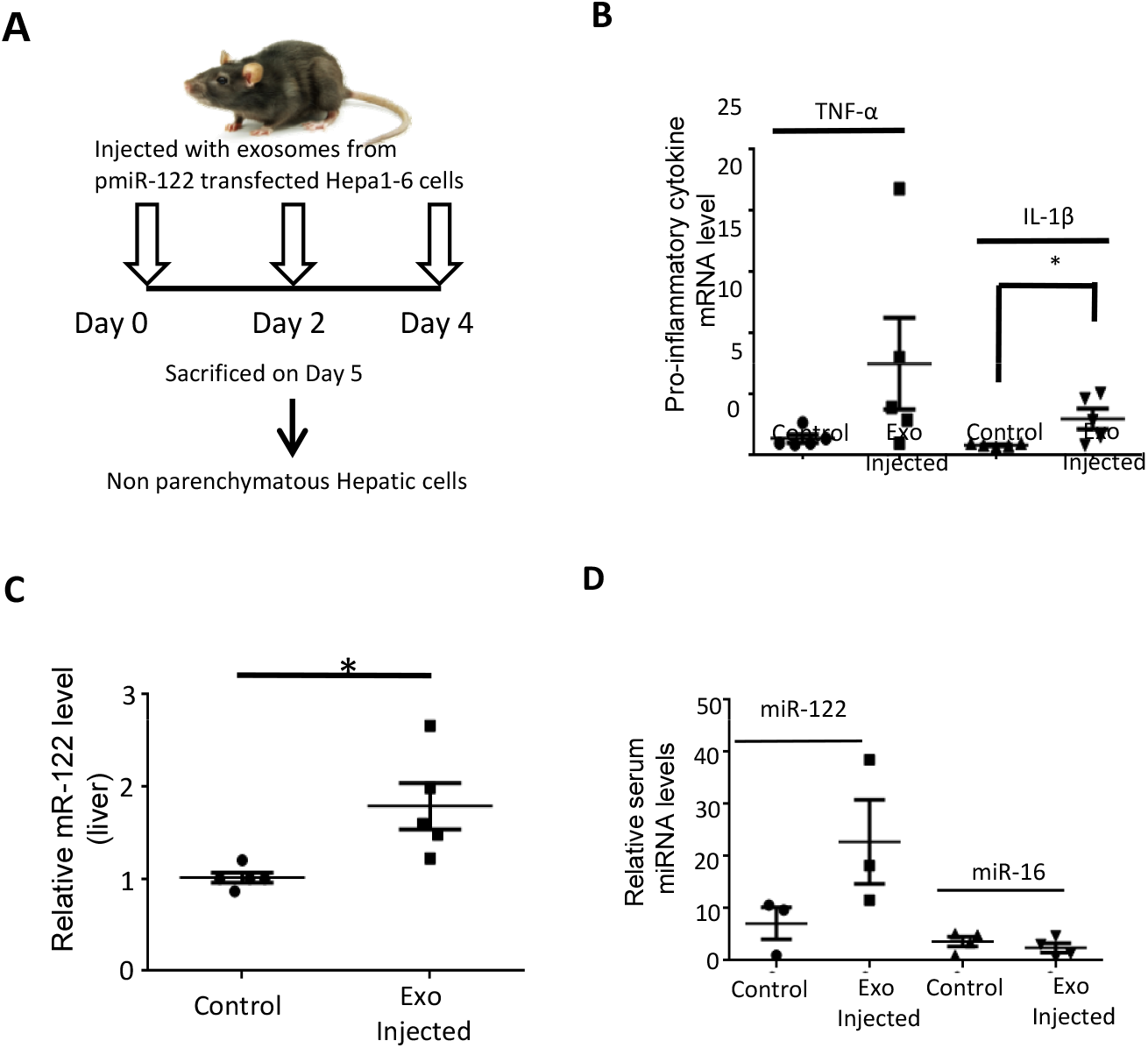
Internalization of external miR-122 by hepatic immune cells *in vivo* elevates expression of proinflammatory cytokine mRNAs. **A-D** Schematic diagram to show the tail vein injection schedule in mice (A). 1x 10^9^ EVs isolated from miR-122 expressing Hepa1-6 cells were injected through the tail vein as per the given schedule for each mice. Equal volume of PBS was administered as the control. Mice were sacrificed on day 5 and non parenchymatous hepatic cells isolated from the liver. Cytokine mRNA (TNF-α and IL-1β) levels in non parenchymatous hepatic cells isolated from EV-injected and control (PBS) injected mice have been shown here (B). Relative internalized miR-122 levels in non parenchymatous hepatic cells are shown (C). Levels of circulatory miR-122 and miR-16 in serum RNA of EV and PBS injected mice are shown (D). All mRNA and miRNA detections were done by RT-qPCR. RT-qPCR detection of cellular miRNAs was done from 25 ng of cellular RNA. RT-qPCR detection of EV miRNAs was done from 100 ng of isolated EV RNA. Normalization of miR-122 and miR-16 were done with respect to U6 snRNA. Normalization of precursor miR-122 and cytokine mRNAs were done with respect to 18S rRNA. Data represents Mean ± SD. (A) N= 3 replicates. (B-D) All data are representative of at least 4 independent mice from each group. For detection of circulatory miRNA (D) serum was isolated from 3 mice of each group. For statistical significance, minimum three independent experiments were considered in each case unless otherwise mentioned and error bars are represented as mean ± S.D. P-values were calculated by utilising Student’s t-test. ns: non-significant, *P < 0.05, **P < 0.01, ***P < 0.0001.

**Supplementary Figure S2.**
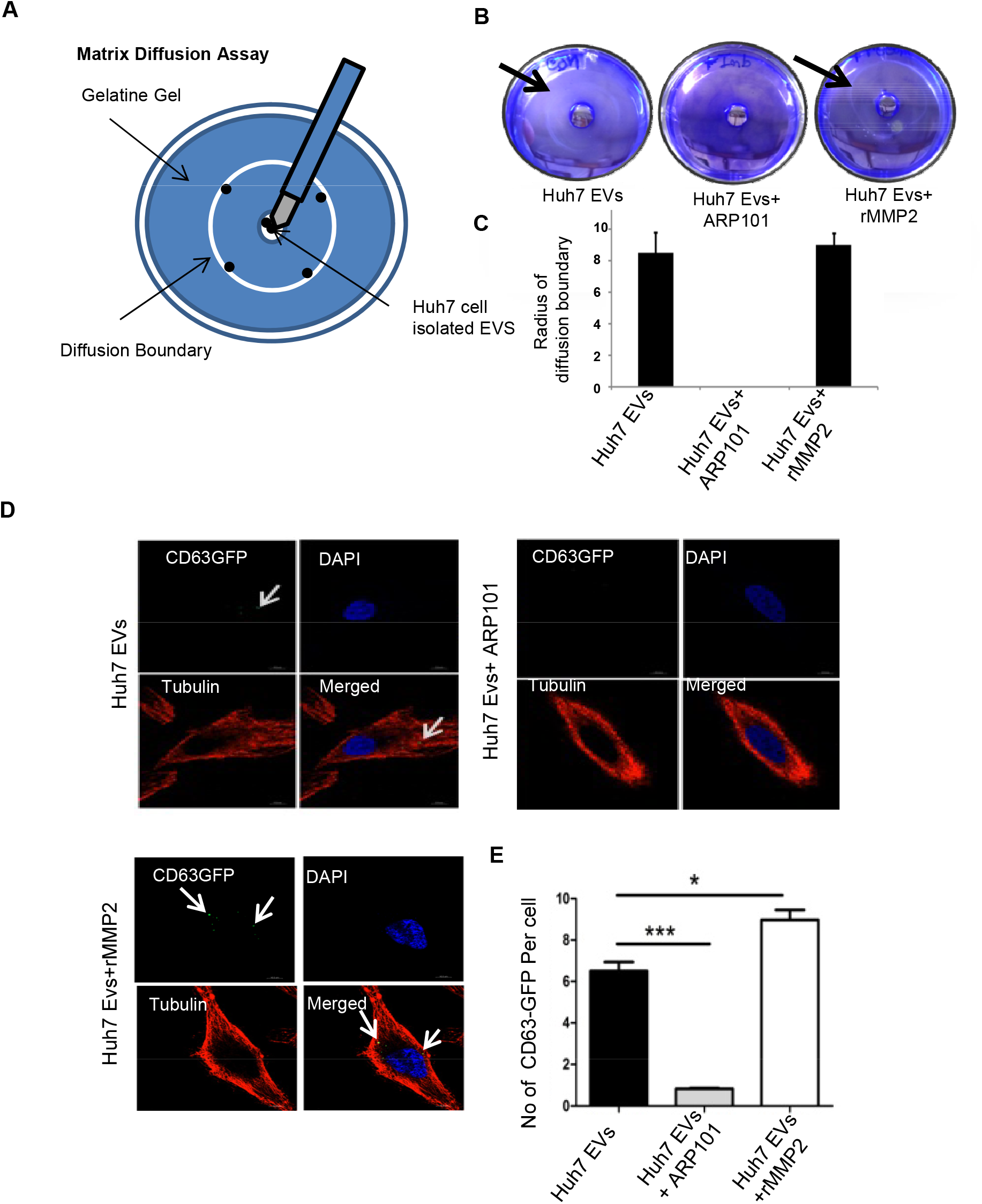
MMP2 facilitates movement of EVs across the extracellular matrix. **A** The diffusion rate measurement assay done with Huh7 released EVs pretreated with rMMP2 or ARP101 as MMP2 inhibitor. EVS were isolated from culture supernatant of Huh7 cells (1×10^7^ Cells). The EVs were then spotted on a well formed in middle of a 35mm Petri disc layered with 2mm thick gelatine matrix. After 30 minutes the reaction were stopped and the gelatin matrix were stained with Brilliant Blue to visualize the diffusion front of the EVs through the matrix. **B-C** The diffusion boundary of EVS in presence and absence of MMP inhibitor ARP101 and rMMP2. The representative plate photographs with marked boundary postions have been shown (B). The diffusion rate calculated from the distance travelled in mm in unit time has been plaotted based on three measurements (n=3) (C). **D-E** MMP2 also facilitate uptake of EVs in recipient cells. The CD63-GFP positive EV transfer from donor cell to recipient HeLa cell grown without the matrigel. The experiments were done in presence and absence of MMP2 inhibitor ARP101 and rMMP2 (D). The quantification were done microscopically and plotted (E) The total number was also quantified by counting green dots of CD63 from three different sets, 5 fields/set, 5 cells/field. For statistical significance, minimum three independent experiments were considered in each case unless otherwise mentioned and error bars are represented as mean ± S.E.M. P-values were calculated by utilising Student’s t-test. ns: non-significant, *P < 0.05, **P < 0.01, ***P < 0.0001.

## References

1. Alisi A, Carpino G, Oliveira FL, Panera N, Nobili V, Gaudio E (2017) The Role of Tissue Macrophage-Mediated Inflammation on NAFLD Pathogenesis and Its Clinical Implications. Mediators of inflammation 2017: 8162421

2. Bartel DP (2009) MicroRNAs: target recognition and regulatory functions. Cell 136: 215–233

3. Bartel DP (2018) Metazoan MicroRNAs. Cell 173: 20–51

4. Basu S, Bhattacharyya SN (2014) Insulin-like growth factor-1 prevents miR-122 production in neighbouring cells to curtail its intercellular transfer to ensure proliferation of human hepatoma cells. Nucleic acids research 42: 7170–7185

5. Becker A, Thakur BK, Weiss JM, Kim HS, Peinado H, Lyden D (2016) Extracellular Vesicles in Cancer: Cell-to-Cell Mediators of Metastasis. Cancer cell 30: 836–848

6. Berg AH, Scherer PE (2005) Adipose tissue, inflammation, and cardiovascular disease. Circulation research 96: 939–949

7. Caballero F, Fernandez A, Matias N, Martinez L, Fucho R, Elena M, Caballeria J, Morales A, Fernandez-Checa JC, Garcia-Ruiz C (2010) Specific contribution of methionine and choline in nutritional nonalcoholic steatohepatitis: impact on mitochondrial S-adenosyl-L-methionine and glutathione. The Journal of biological chemistry 285: 18528–18536

8. Cai D, Yuan M, Frantz DF, Melendez PA, Hansen L, Lee J, Shoelson SE (2005) Local and systemic insulin resistance resulting from hepatic activation of IKK-beta and NF-kappaB. Nature medicine 11: 183–190

9. Chang J, Nicolas E, Marks D, Sander C, Lerro A, Buendia MA, Xu C, Mason WS, Moloshok T, Bort R, Zaret KS, Taylor JM (2004) miR-122, a mammalian liver-specific microRNA, is processed from hcr mRNA and may downregulate the high affinity cationic amino acid transporter CAT-1. RNA biology 1: 106–113

10. Chevillet JR, Kang Q, Ruf IK, Briggs HA, Vojtech LN, Hughes SM, Cheng HH, Arroyo JD, Meredith EK, Gallichotte EN, Pogosova-Agadjanyan EL, Morrissey C, Stirewalt DL, Hladik F, Yu EY, Higano CS, Tewari M (2014) Quantitative and stoichiometric analysis of the microRNA content of exosomes. Proceedings of the National Academy of Sciences of the United States of America 111: 14888–14893

11. Esau C, Davis S, Murray SF, Yu XX, Pandey SK, Pear M, Watts L, Booten SL, Graham M, McKay R, Subramaniam A, Propp S, Lollo BA, Freier S, Bennett CF, Bhanot S, Monia BP (2006) miR-122 regulation of lipid metabolism revealed by in vivo antisense targeting. Cell metabolism 3: 87–98

12. Filipowicz W, Bhattacharyya SN, Sonenberg N (2008) Mechanisms of post-transcriptional regulation by microRNAs: are the answers in sight? Nature reviews Genetics 9: 102–114

13. Han KY, Dugas-Ford J, Seiki M, Chang JH, Azar DT (2015) Evidence for the Involvement of MMP14 in MMP2 Processing and Recruitment in Exosomes of Corneal Fibroblasts. Investigative ophthalmology & visual science 56: 5323–5329

14. Han L, Sheng B, Zeng Q, Yao W, Jiang Q (2020) Correlation between MMP2 expression in lung cancer tissues and clinical parameters: a retrospective clinical analysis. BMC pulmonary medicine 20: 283

15. Jo YK, Park SJ, Shin JH, Kim Y, Hwang JJ, Cho DH, Kim JC (2011) ARP101, a selective MMP-2 inhibitor, induces autophagy-associated cell death in cancer cells. Biochemical and biophysical research communications 404: 1039–1043

16. Jopling C (2012) Liver-specific microRNA-122: Biogenesis and function. RNA biology 9: 137–142

17. Lotvall J, Valadi H (2007) Cell to cell signalling via exosomes through esRNA. Cell Adh Migr 1: 156–158

18. Momen-Heravi F, Bala S, Kodys K, Szabo G (2015) Exosomes derived from alcohol-treated hepatocytes horizontally transfer liver specific miRNA-122 and sensitize monocytes to LPS. Scientific reports 5: 9991

19. Morinaga H, Mayoral R, Heinrichsdorff J, Osborn O, Franck N, Hah N, Walenta E, Bandyopadhyay G, Pessentheiner AR, Chi TJ, Chung H, Bogner-Strauss JG, Evans RM, Olefsky JM, Oh DY (2015) Characterization of distinct subpopulations of hepatic macrophages in HFD/obese mice. Diabetes 64: 1120–1130

20. Mukherjee K, Ghoshal B, Ghosh S, Chakrabarty Y, Shwetha S, Das S, Bhattacharyya SN (2016) Reversible HuR-microRNA binding controls extracellular export of miR-122 and augments stress response. EMBO reports 17: 1184–1203

21. Pirola CJ, Fernandez Gianotti T, Castano GO, Mallardi P, San Martino J, Mora Gonzalez Lopez Ledesma M, Flichman D, Mirshahi F, Sanyal AJ, Sookoian S (2015) Circulating microRNA signature in non-alcoholic fatty liver disease: from serum non-coding RNAs to liver histology and disease pathogenesis. Gut 64: 800–812

22. Shimoda M, Khokha R (2017) Metalloproteinases in extracellular vesicles. Biochimica et biophysica acta Molecular cell research 1864: 1989–2000

23. Shoelson SE, Lee J, Goldfine AB (2006) Inflammation and insulin resistance. The Journal of clinical investigation 116: 1793–1801

24. Tan Y, Ge G, Pan T, Wen D, Gan J (2014) A pilot study of serum microRNAs panel as potential biomarkers for diagnosis of nonalcoholic fatty liver disease. PloS one 9: e105192

25. Tang H, He Y, Li L, Mao W, Chen X, Ni H, Dong Y, Lyu F (2019) Exosomal MMP2 derived from mature osteoblasts promotes angiogenesis of endothelial cells via VEGF/Erk1/2 signaling pathway. Experimental cell research 383: 111541

26. Valadi H, Ekstrom K, Bossios A, Sjostrand M, Lee JJ, Lotvall JO (2007) Exosome-mediated transfer of mRNAs and microRNAs is a novel mechanism of genetic exchange between cells. Nature cell biology 9: 654–659

27. Wang B, Ding YM, Fan P, Xu JH, Wang WX (2014) Expression and significance of MMP2 and HIF-1alpha in hepatocellular carcinoma. Oncology letters 8: 539–546

28. Wang Y, Liang H, Jin F, Yan X, Xu G, Hu H, Liang G, Zhan S, Hu X, Zhao Q, Liu Y, Jiang ZY, Zhang CY, Chen X, Zen K (2019) Injured liver-released miRNA-122 elicits acute pulmonary inflammation via activating alveolar macrophage TLR7 signaling pathway. Proceedings of the National Academy of Sciences of the United States of America 116: 6162–6171

